# What causes bird-building collision risk? Seasonal dynamics and weather drivers

**DOI:** 10.1101/2022.09.30.510341

**Authors:** Kara M. Scott, Attilla Danko, Paloma Plant, Roslyn Dakin

## Abstract

Bird-building collisions are a major source of wild bird mortality, with hundreds of millions of fatalities each year in the United States and Canada alone. Here, we use two decades of daily citizen-science monitoring to characterize day-to-day variation in building collisions and determine the factors that predict the highest risk times in two North American cities. We use these analyses to evaluate three potential causes of increased collision risk: heightened migration traffic during benign weather, increased navigational and flight errors during inclement weather, and increased errors in response to highly directional sunlight that enhances reflected images. The seasonal phenology of collisions was consistent across sites and years, with daily collision rates approximately two-fold higher in autumn as compared to spring. During both migration seasons, collision risk was best predicted by the weather conditions at dawn. In spring, peak collision risk occurs on days with warm temperatures, south winds, and a lack of precipitation at dawn. In autumn, peak collision occurs on days with cool temperatures, north winds, high atmospheric pressure, a lack of precipitation, and clear conditions with high visibility. Based on these results, we hypothesize that collisions are influenced by two main weather-driven mechanisms. First, benign weather at dawn with winds that are favorable for migration flight causes an increase in migration traffic in both spring and autumn, creating greater opportunity for collisions to occur. Second, for autumnal migrants, cold clear conditions may cause an additional increase in collision risk. We propose that these conditions may be particularly hazardous in autumn because of the high abundance of naïve and diurnal migrants at that time of year. Our analysis also establishes that a relatively small proportion of days (15%) are responsible for 50% of the total collision mortality within a season, highlighting the importance of targeting mitigation strategies to the most hazardous times.

## 1. Introduction

Birds have been in a marked decline, and North America has lost an approximate 29% of bird abundance since the 1970s (Rosenberg et al., 2019). One factor that may contribute to these losses is the widespread mortality caused by birds colliding with buildings and windows, which are largely driven by collisions by migrating birds (Arnold & Zink, 2011; Elmore et al., 2021; Loss, Will, Loss, & Marra, 2014; Loss, Will, & Marra, 2015). Indeed, meta-analyses have estimated that 365–988 million birds are killed in collisions each year in the USA, while another 16–42 million are killed in collisions each year in Canada (Loss et al., 2014; Machtans et al., 2013). Collision risk varies greatly across landscapes, buildings, seasons, days, and times of day. Given that collisions occur during flight, weather may be an important driver of this variation because atmospheric conditions influence migration decisions, flight and navigational errors, and the visual appearance of glass. Understanding the temporal dynamics of these collisions is important for two reasons: it allows us to better predict times that will have elevated risk, and it provides insight into potential causal mechanisms that can inform mitigation. This knowledge is important for aeroecology and aeroconservation, which are underrepresented areas of study (Davy, Ford, & Fraser, 2017; Horton, Van Doren, Stepanian, Farnsworth, & Kelly, 2016).

Several previous studies have examined the question of how weather conditions may predict bird collisions. An early study by Klem (1989) reported a sample of anecdotal descriptions of weather conditions at 2 houses on days when 83 collisions occurred, and hypothesized that collisions at houses may be higher during benign weather. A 2020 study by Loss et al. examined the weather conditions associated with collisions of 15 individual American Woodcocks at 21 houses during spring 2018, and found that these Woodcock collisions were associated with inclement weather events involving strong north winds (which opposed the direction of their pre-breeding migration at that time) (Loss et al., 2020). While these observations are valuable, they do not allow us to draw generalized conclusions regarding collision dynamics. Most recently, Lao et al. (2022) analyzed daily data on several hundred collisions that occurred at 21 buildings in Minneapolis, and reported that collision risk was greater during conditions with wind directions that were favorable for migration, as well as with changing weather patterns. Although Lao’s (2022) study is the most rigorous to date, it is still limited geographically. Here, we leverage information on more than 62,000 migratory bird collisions, representing two decades of daily monitoring by citizen scientists in two large metropolitan areas (Toronto, Canada, and Chicago, USA). Our goal is to evaluate the influence of weather and atmospheric conditions on daily variation in avian collision rates in these urban environments.

There are several non-exclusive mechanisms by which weather and atmospheric conditions might influence collision risk for birds. The first possibility is that weather-induced changes in migration traffic (defined as the abundance of migrating birds in the air) can influence the occurrence of collisions. Consistent with this hypothesis, avian collision rates with buildings and aircraft broadly correlate with times of higher migration traffic (Elmore et al., 2021; Lao et al., 2022; Nilsson et al., 2021; van Doren et al., 2021). Increased migration traffic is commonly driven by benign weather during both spring and autumn migration periods (Bozó, Csörgõ, & Heim, 2018; Erni, Liechti, Underhill, & Bruderer, 2002; Panuccio, Dell’Omo, Bogliani, Catoni, & Sapir, 2019; Welcker & Vilela, 2019). During spring, the strongest predictor of migration traffic is often warmer temperatures (Bozó et al., 2018; Haest, Hüppop, & Bairlein, 2020; Hamer et al., 2021; Schekler & Sapir, 2021; Van Doren & Horton, 2018; Welcker & Vilela, 2019). By contrast, in autumn, the effects of temperature on migration traffic are less clear, with different studies finding positive, negative, or no associations between temperatures and autumn migration traffic (Bozó et al., 2018; Erni et al., 2002; Loss et al., 2020; Nilsson et al., 2019; Schekler & Sapir, 2021; Welcker & Vilela, 2019). If weather-induced changes in migration traffic are a major driver of avian collisions, then we expect higher collision rates to occur on days with benign conditions in both spring and fall (i.e., favorable wind direction, low wind speeds, low precipitation, and high atmospheric pressure), with additional risk posed by warm temperatures in spring.

A second mechanism by which weather and atmospheric conditions might influence collision risk is that inclement weather may increase the probability that birds fail to avoid collisions during flight. For example, inclement weather may negatively impact visual guidance and the ability of birds to detect and respond appropriately to obstacles (Marques et al., 2014). Inclement weather could also increase physiological stress through depletion of energy reserves, making birds more likely to make fatal errors. This is occasionally observed in mass casualty events; for example, in the fall of 2020, winter storm conditions led to a mass bird casualty event in the southwestern USA (Johnson, 2020). Certain weather conditions may also cause flying migrants to descend to lower altitudes, where the risk of collisions is greater. For instance, it has been proposed that inclement weather and cloud cover may drive flying birds down to lower altitudes that have more artificial light at night (ALAN) and more obstacles (Cochran & Graber, 1958; Van Doren et al., 2021). In general, if weather-induced flight and navigational errors are a major driver of collisions, then we then expect higher collision rates to occur on days with high wind speeds, high precipitation, increased cloud cover, and cold temperatures, all of which may disrupt sensory performance, increase stress, and/or drive flying birds to change flight altitude.

A third mechanism by which atmospheric conditions might influence collision risk is by affecting the visual appearance of glass windows. The appearance of glass has been found to be an important proximate driver of collisions, based on positive associations between collision rates and the extent of glass on buildings (Riding, O’Connell, & Loss, 2020), and experiments demonstrating that visual deterrents and acoustic signals placed on or in front of windows can prevent collisions (Boycott, Mullis, Jackson, & Swaddle, 2021; Klem, 2009; Klem & Saenger, 2013). The appearance of glass is also strongly influenced by weather: on days with clear skies (i.e., little to no cloud cover and/or high visibility), sunlight is more directional, which creates specular reflections and brightly-coloured reflected images of the surroundings on glass. These clear-sky conditions may increase the probability that birds are deceived by reflected images. By contrast, on cloudy or overcast days, and on days with precipitation, sunlight is diffuse, and the brightness and coherence of reflections is reduced due to low coherence as well as water droplets on reflective surfaces. If weather-induced changes in the appearance of glass are an important driver of collisions, then we expect collision rates to be elevated on days with reduced cloud cover, high atmospheric visibility, and a lack of precipitation.

In addition to the mechanisms above, we also consider the possibility that the phase of the moon may influence daily variation in collision risk (Norevik, Åkesson, Andersson, Bäckman, & Hedenström, 2019). The moon can be a navigational beacon for some nocturnal migrants and may facilitate flight at night (Miles, Money, Luxmoore, & Furness, 2010; Syposz, Gonçalves, Carty, Hoppitt, & Manco, 2018). In addition to its potential effects on migration traffic, it is also possible that lunar illumination may interact with ALAN and/or other sources of navigational disruption that may influence collision risk. We therefore include lunar illumination in our analysis, although we have no *a priori* predictions as to the direction of any potential lunar effects.

Here, we investigate the weather conditions that predict day-to-day variation in building collisions for a broad range of North American migratory birds. Our analysis considers nine variables: air temperature, wind speed, wind direction, precipitation, relative humidity, atmospheric pressure, cloud cover, atmospheric visibility, and lunar illumination. We examine these potential drivers using data on over 62,000 collisions involving more than 100 migratory bird species. Our goal is to identify the general drivers of collisions in urban environments, rather than species-specific effects. We first use these data to estimate day-to-day variation in collision rates, and compare spring and autumn seasons. Second, we quantify the extent to which collisions are concentrated over a small number of dates (Horton, Van Doren, Albers, Farnsworth, & Sheldon, 2021). Third, we evaluate if daily collision rates are best explained by weather conditions at a specific time of day (morning, dawn, overnight, or the previous evening). Finally, we describe the relationships between collision rates and weather during spring and autumn, respectively. Our overall aim is to identify the major drivers of collision risk as a first step to identifying causal mechanisms. Ultimately, with increased knowledge, improved mitigation and aeroconservation measures will be possible (Davy et al., 2017).

## 2. Methods

### 2.1 Collision rates

To estimate daily collision rates, we obtained data on the number of migratory bird collision casualties recorded per day from two large metropolitan areas: Toronto, Canada (43.74 N, 79.37 W) and Chicago, USA (41.88 N, 87.63 W). Both cities have a long-term daily collision monitoring programs that provide the basis for important continent-wide estimates of the extent and causes of collision mortality (Arnold & Zink, 2011; Colling, Guglielmo, Bonner, & Morbey, 2022; Cusa, Jackson, & Mesure, 2015; Evans Ogden, 1996; Loss et al., 2014; Machtans et al., 2013; Van Doren et al., 2021; Winger et al., 2019). The data from Toronto were collected by FLAP Canada (Fatal Light Awareness Program, www.flap.org) and in Chicago by the Chicago Bird Collision Monitors and McCormick Place monitors (www.birdmonitors.net) (Winger et al., 2019).

For the purpose of this study, we focused on a 10-year period from each city with daily records on the number of migratory bird collision casualties detected (years 2009 to 2018 in Toronto, and 2006 to 2015 in Chicago). We chose these particular years because they represent the contiguous 10-year time span within each city that had the largest number of collision records available. The total number of records in the data included 28,657 migratory bird casualties in Toronto and 33,493 in Chicago. The Chicago dataset focused on casualties from the passerine order (89 identified species), whereas the Toronto dataset was predominantly passerines, with some non-passerines also recorded (138 identified bird species, 92% of which were passerines). All data processing and analyses were performed in R 4.2.2.

The vast majority of building collisions occur during the periods from March–June, and August– November, representing spring and autumn migration seasons, respectively. During the months of July, December, January, and February, over 95% of dates had 0 collision records in our data. Moreover, only 0.2% of all collision records were attributed to dates within the predominantly non-migratory months. Given that our aim in this study was to evaluate sources of day-to-day variation in avian collision rates, we limited our analyses to the spring and autumn migration seasons of March–June and August–November, as day-to-day variation was not present at other times (Fig. 1). The total number of city-dates in our analysis was 1,465 days in spring (745 from Toronto and 720 from Chicago) and 1,855 in autumn (925 from Toronto and 930 from Chicago). For the 1,023 dates that overlapped between the two cities, daily log-counts of collisions were moderately correlated (Pearson’s R = 0.52, p < 0.0001).

**Figure 1.**
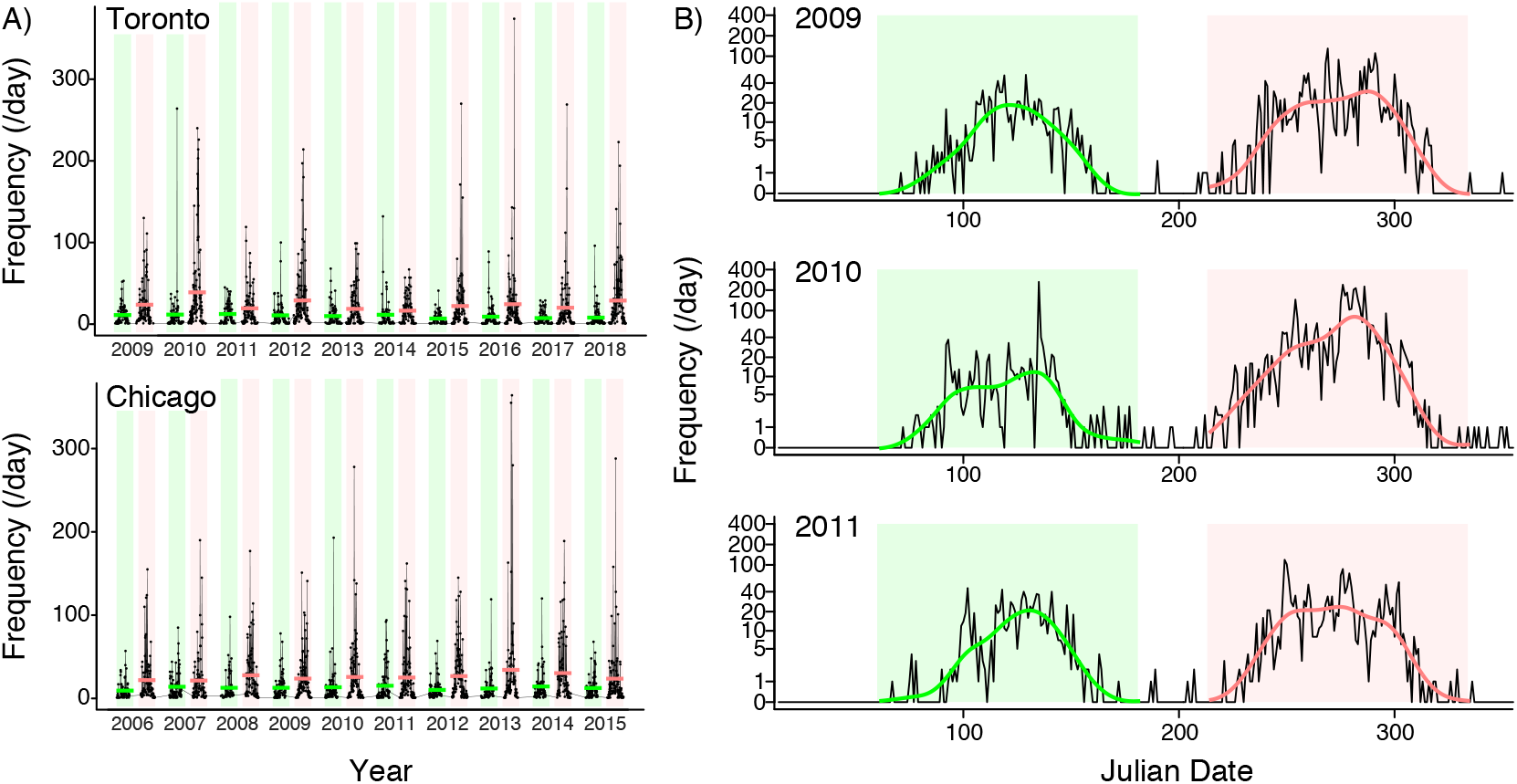
Seasonal phenology of bird collisions with buildings. (A) Daily collision rates recorded in two cities, Toronto and Chicago, during the spring (green) and autumn (red) migration periods (horizontal lines show the means). Daily collision rates are consistently higher during autumn migration, as compared to spring. Note that the span of studied years differs for the two sites. (B) To evaluate atmospheric and weather drivers of collision rates while controlling for seasonal phenology, we first determined the expected rate of collisions by fitting a cubic spline to each seasonal peak (shown by the green and pink lines in B). This procedure was repeated for each site-season dataset; examples from Toronto are shown here. Next, we analyzed the daily collision rate while controlling for the expected phenological pulse within each season. Hence, the analysis investigates how weather conditions may explain fluctuations above or below the seasonal expected rate of collisions.

### 2.2 Weather and atmospheric conditions

We obtained hourly weather station data on nine weather and atmospheric parameters that are expected to influence flight and collision risk: air temperature (C), wind speed (km/h), wind direction (°), precipitation (mm/h), relative humidity (%), atmospheric pressure (kPa), cloud cover (%), atmospheric visibility (km), and lunar illumination (%). Hourly values were obtained for each city from www.openweathermap.org. All cloud cover estimates were expressed as a percent of full cloud cover, such that a value of 100% indicates a completely overcast sky. Lunar illumination values represent a measure of lunar phase and were obtained from the R package lunar 0.1-04; a value of 0% indicates the new moon and 100% indicates a full moon.

We assume that collisions recorded on a given day may have occurred at any time in the previous 24 hours, with weather during the previous 12-14 hours hypothesized to have the greatest influence (Evans Ogden, 1996; Lao et al., 2020). Therefore, we summarized the weather data into four time periods: (i) the previous evening, defined as the average of hourly weather conditions from 2000 to 2300 on the previous day; (ii) the overnight period, defined as the average of hourly conditions from 0000 to 0300; (iii) dawn, defined as the average of hourly conditions from 0400 to 0700; and (iv) morning, defined as the average of hourly conditions from 0800 to 1100 (all local times). As an example, for collisions recorded on April 10, 2009, the corresponding local morning temperature would be calculated as the average from 0800-1100 that day, whereas the corresponding previous evening temperature would be calculated as the average from 2000-2300 on April 9, 2009.

### 2.3 Migration collision phenology

Collision rates have a well-defined phenological peak within each spring and autumn migration season, corresponding with avian migration phenology (Fig. 1). To quantify this baseline temporal variation in migration collision phenology, we fit a cubic spline to the daily number of casualties as a function of day of the year within each city-year-season, with 0.6 as the smoothing parameter (Fig. 1B). The resulting splines represent the expected phenological pulse of casualties for a given season and year, at a given location. We used these phenology pulse estimates as a covariate in our statistical analysis of day-to-day variation in collision rates (see details in section 2.4 below). Hence, our analysis evaluates if local daily weather can explain day-to-day variation in collision rates, after accounting for the expected seasonal pulse due to the timing of migration in a given year (Fig. 1B).

### 2.4 Statistical analysis of collision rates and weather

We used mixed-effects regression models to evaluate sources of variation in daily collision rates. The response variable was the daily number of collisions recorded (per city-date), log-transformed to meet the assumptions of linear regression. All models were fit in R using the lme4 1.1-30 and lmerTest 3.1-3 packages.

To provide descriptive statistics for seasonal differences in collision rates, we first fit a regression model with season as the only fixed effect predictor (spring or autumn), and year and city as random effects.

Next, we evaluated which weather time period was the best predictor of daily collision rates within each season. To do this, we compared five candidate mixed-effect regression models: a previous evening model, an overnight model, a dawn model, a morning model, and a null model. All models included migration collision phenology as a fixed effect predictor. The time period models included as additional fixed effects the nine weather variables described in section 2.2 above. Hence, a morning model contained local weather conditions for the morning time period, whereas a dawn model contained local weather conditions for the dawn period, etc. Because wind direction is a circular variable (360° = 0°), we modelled wind direction as a 2^nd^-order orthogonal polynomial. This was important because it allows wind directions at the boundary (0° and 360°) to have equivalent effects. In the context our study sites in Toronto and Chicago, south winds are expected to be favorable in spring, and north winds in autumn. The null model included migration collision phenology as a fixed effect predictor, but had no weather variables. In all models, year and city were included as random effects.

Regression models were fit with maximum likelihood and compared using Akaike’s Information Criterion (AIC). We also used the MuMIn package 1.46.0 to calculate Akaike weights. The Akaike weight is a measure of the relative strength of the evidence for a given model in a model set, expressed as a probability ranging from 0 to 1; the weights for all models in a set sum to 1. Hence, if a model has an Akaike weight of 1, it indicates complete certainty that the model is best supported within its model set. If two models receive similar Akaike weights, it indicates that they are equivalently well-supported. We performed this model selection analysis separately for each season (spring and autumn).

Finally, we examined the coefficient estimates from the best-supported regression model for spring and autumn, respectively. We used mean-centred and standardized variables (response and predictor) to allow comparison of effect sizes within and across models. We also obtained p-values using Satterthwaite’s method in the lmerTest package. As a measure of the total variance in collision rates explained by the fixed effects in each model, we calculated R^2^_GLMM_ using the MuMIn package. As an additional check on our results, we verified that day of the week was not a significant predictor of collision rates in our analyses, and we confirmed that all conclusions were unchanged when controlling for day of the week.

One possibility that we do not consider here is the potential that collision risk could be influenced by interactive effects of several weather variables (wherein one weather parameter modulates the effect others). Similarly, we do not consider combinations of weather conditions from different time periods in the present study. Instead, for simplicity and greatest generalizability, we limit our analyses here to only considering the main effects of weather variables within a given time period.

## 3. Results

### 3.1 Seasonal and day-to-day variation in collision rates

The average daily collision rate was approximately twice as high in autumn as compared to spring (1.86-fold greater in autumn; p < 0.0001; Fig. 1A). Within each season, there was substantial day-to-day variation in collision rates (Fig. 1B). Notably, a relatively small number of days account for a large proportion of the seasonal total (Fig. 2). For example, the 12 worst days typically account for about 50% of the seasonal total, and the three worst days typically account for between 19-22% of the seasonal total. In spring, collision risk consistently peaked in the 17^th^ week of the year (the last days of April and first days of May). In autumn, collisions consistently peaked in the 40^th^ week of the year (the first week of October).

**Figure 2.**
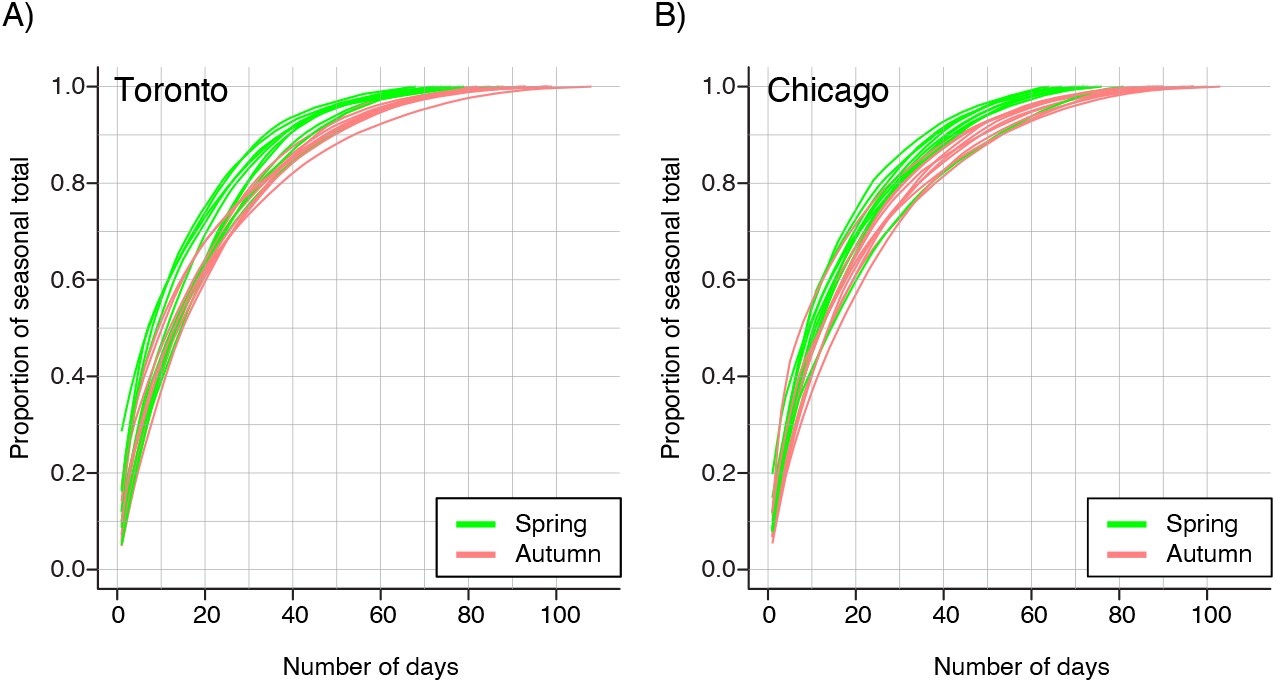
Most collision risk is concentrated within a small number of days. These plots show the cumulative proportion of total seasonal mortality for (A) Toronto and (B) Chicago. To perform this analysis, days were first ranked by severity within each site-season (1^st^ = most severe, etc.). Then, each day’s cumulative proportion of the seasonal total was plotted with lines connecting data from a given site-season. The steep increase in these cumulative proportion curves illustrates how most collisions can be attributed to a small number of severe days within a season. Each season represents four months: spring = March through June, and autumn = August to November.

### 3.2 Drivers of spring collisions

Dawn weather was the best predictor of spring collision risk (Akaike weight > 0.57; Table S1). The null model, which included phenology, but no weather effects, had an Akaike weight of 0. The best-supported model using weather at dawn indicates that spring collisions are mainly driven by warm temperatures, south winds (which are favorable for north-flying migrants), and dry conditions with a lack of precipitation (Fig. 3A and Table 1).

**Figure 3.**
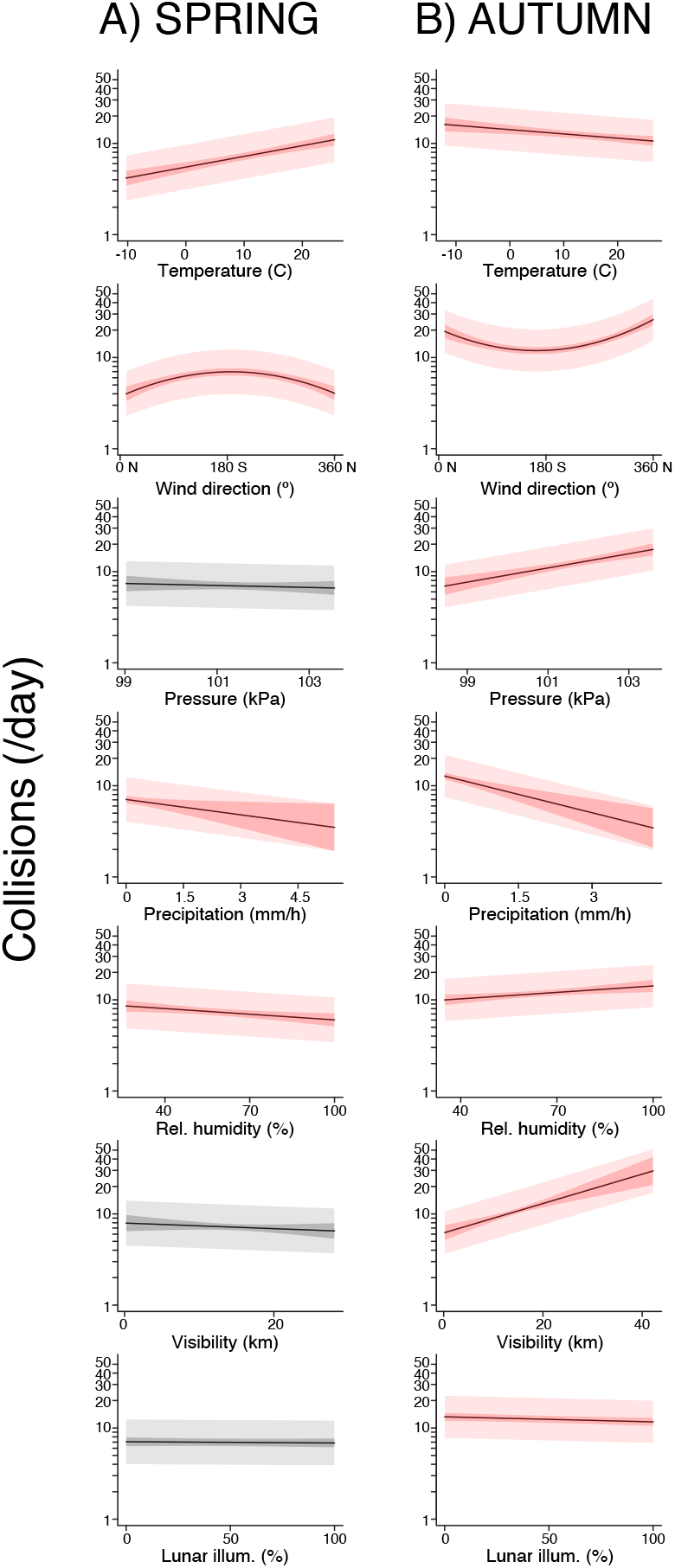
Collision risk is driven by weather conditions that are largely benign for flight, as well as high-visibility days in autumn. (A) In spring, high collision rates are associated with warm temperatures, south (favorable) winds, a lack of precipitation and low humidity. (B) In autumn, high collision rates are associated with cool temperatures, north (favorable) winds, high pressure, a lack of precipitation, and high atmospheric visibility. Each panel in (A) and (B) shows the predictions of the best-supported model when all other predictors are set to the mean value. The inner shaded region shows the 95% CI for the line-of-best fit. The outer shaded region shows the 50% prediction interval of the model. Red is used for predictors that had a statistically significant association with collision rates in the full model. For simplicity, only the weather parameters that explain a substantial amount of collision variation in at least one season are presented in this plot. See Tables 1-2 and Tables S1-S2 for additional details of this analysis.

**Table 1.** Weather drivers of bird-building collisions in spring. The spring weather conditions associated with the greatest collision risk include warm temperatures with a favorable (south) wind, a lack of precipitation, and low relative humidity. To facilitate comparison of effect sizes, the response variable and all predictor variables were centered and standardized to have mean = 0 and SD = 1. Predictors are indicated in bold where the slope is statistically significant. See Tables S1 for additional details.

| Predictor | Estimate | 95% CI | p |
| --- | --- | --- | --- |
| **Temperature** | **0.14** | **[0.10, 0.18]** | **< 0.0001** |
| Wind speed | -0.02 | [-0.07, 0.02] | 0.27 |
| Poly(Wind direction)-1 | -0.56 | [-2.13, -1.01] | 0.48 |
| **Poly(Wind direction)-2** | **-4.90** | **[-6.40, -3.40]** | **< 0.0001** |
| **Precipitation** | **-0.05** | **[-0.09, -0.01]** | **0.02** |
| **Relative humidity** | **-0.06** | **[-0.11, -0.02]** | **0.002** |
| Atmos. Pressure | -0.02 | [-0.06, 0.03] | 0.48 |
| Cloud cover | 0.04 | [0.00, 0.09] | 0.052 |
| Visibility | -0.03 | [-0.07, 0.01] | 0.11 |
| Lunar illum. | -0.01 | [-0.05, 0.03] | 0.62 |
| **Phenology** | **0.62** | **[0.58, 0.66]** | **< 0.0001** |

### 3.3 Drivers of autumn collisions

Dawn weather was also the best predictor of autumn collision rates (Akaike weight > 0.99; Table S2). The null model for autumn had an Akaike weight of 0. The best-supported model using weather at dawn indicates that autumn collisions are mainly driven by cool temperatures, north winds (which are favorable for south-flying migrants), a lack of precipitation, high relative humidity, high atmospheric pressure, and high visibility (Fig. 3B and Table 2).

**Table 2.** Weather drivers of bird-building collisions in autumn. The autumn weather conditions associated with the greatest collision risk include cold temperatures with a favorable (north) wind, a lack of precipitation, high visibility, and high atmospheric pressure. To facilitate comparison of effect sizes, the response variable and all predictor variables were centered and standardized to have mean = 0 and SD = 1. Predictors are indicated in bold where the slope is statistically significant. See Tables S2 for additional details.

| Predictor | Estimate | 95% CI | p |
| --- | --- | --- | --- |
| **Temperature** | **-0.05** | **[-0.08, -0.02]** | **0.0006** |
| Wind speed | -0.03 | [-0.06, 0.00] | 0.06 |
| **Poly(Wind direction)-1** | **3.42** | **[2.17, 4.69]** | **< 0.0001** |
| **Poly(Wind direction)-2** | **5.29** | **[4.06, 6.51]** | **< 0.0001** |
| **Precipitation** | **-0.08** | **[-0.10, -0.05]** | **< 0.0001** |
| **Relative humidity** | **0.05** | **[0.01, 0.09]** | **0.006** |
| **Atmos. Pressure** | **0.10** | **[0.06, 0.13]** | **< 0.0001** |
| Cloud cover | -0.02 | [-0.05, 0.01] | 0.16 |
| **Visibility** | **0.14** | **[0.09, 0.18]** | **< 0.0001** |
| **Lunar illum.** | **-0.03** | **[-0.06, -0.01]** | **0.01** |
| **Phenology** | **0.78** | **[0.75, 0.81]** | **< 0.0001** |

There was also a tendency for elevated collision rates after dark autumn nights with low lunar illumination (Table 2). However, as shown in Fig. 3B, this lunar illumination effect does not explain as much of the day-to-day variation in collision rates as the effects of temperature, wind, precipitation/pressure and visibility.

## 4. Discussion

We show here that bird building collisions follow a consistent seasonal phenology with strong peaks in the spring and autumn migration seasons (Fig. 1). Our analysis also shows that autumn migration is particularly hazardous (Fig. 1A), with daily collisions rates approximately twice as severe as spring. The increased risk of collision mortality in autumn is likely due to the presence of numerous first-time, naïve migrants in autumn (Newton, 2007). Although post-breeding autumn migration has cumulatively more individual migrants, studies of bird migration intensity have shown that spring migration generally has greater daily peak levels of migration traffic (i.e., in spring, migration traffic is concentrated with higher peak traffic per day) (Dokter et al., 2018; Ng et al., 2022). The idea that greater daily collision rates in autumn are mainly driven by first-time migrants is also corroborated by recent studies finding that local birds (with greater experience of the landscape) are better at avoiding collisions (Riding et al., 2021; Sabo, Hagemeyer, Lahey, & Walters, 2016).

We also found that a relatively small number of days are responsible for a large burden of total collision mortality within a site. For example, 50% of the recorded collisions occur in just 15% of days within a season (Fig. 2). This finding is consistent with previous studies monitoring the abundance of migrating birds, which have revealed that a relatively small number of days accounts for the majority of individual movements (Horton et al., 2021; Van Doren & Horton, 2018). This concentration of migration (and collisions) into peak days emphasizes the importance of understanding drivers of day-to-day variation.

During spring, we found that day-to-day variation in collision rates was best explained by weather conditions at dawn, consistent with recent reports that collisions peak overnight and during early morning hours (Colling et al., 2022; Riding et al., 2021; Van Doren et al., 2021). Specifically, we found that spring collision risk was greatly elevated on days with warmer temperatures, a lack of precipitation, and south winds (which would be favorable for prebreeding migration at our study sites; Fig. 3A). Based on these results in the context of migration biology, we suggest that the biggest driver of spring collisions is weather-induced changes in migration traffic (Elmore et al., 2021; Hamer et al., 2021). On warm, dry spring days with favorable winds, there is an increased abundance of birds in the air, and this increases the opportunity for collisions to occur.

During autumn, collision rates were also best predicted by the weather at dawn. Specifically, autumn collision mortality was elevated on days that were cold, with high atmospheric pressure, a lack of precipitation, north winds (favorable for post-breeding migration), and high atmospheric visibility (Fig. 3B). These results suggest that autumn collisions may be driven by a combination of two major mechanisms: an increase in migration traffic at dawn during mornings with favorable flight conditions, and an increase in potential errors in response to glass under conditions with highly directional sunlight. We found a weak association between collision rates and times with a darkened moon phase in autumn, but lunar illumination explained much less of the day-to-day variation as compared to other predictors (Fig. 3B).

We designed our analysis to focus on the most generalizable weather predictors for the two broad, urban sites in our data. It is important to note that there may be differences across sites, and for other locations, additional site-specific predictors are possible depending on geography, and the geometry of specific landscape features used by birds as well as that of the buildings themselves. For example, Van Doren et al (2021) found that Lake Michigan factored into weather-collision interactions, where winds could cause higher avian concentrations along Chicago’s shoreline. Inclement weather can also drive mass casualty events that are specific to certain locations (Johnson, 2020; Newton, 2007) and/or species (Loss et al., 2020). It is important to note that even though inclement weather may not be associated with most collisions at our study site, it can still cause many collisions and may even be a contributing factor to collisions that occur on benign days. For example, inclement weather may cause birds to make suboptimal choices for stopover landing areas, e.g., in urban regions or other areas that pose greater mortality risk (Jenni & Schaub, 2003). This may indirectly influence collision rates if birds later attempt to leave the urban area at the next benign weather opportunity (Clipp et al., 2020; Van Doren et al., 2017). This could be due to the greater number of birds forced to initiate flights in a hazardous environment (Lao et al., 2022). It is also possible that ALAN may compound the effects of inclement weather, as both may be disorienting factors (Arnold & Zink, 2011; Evans Ogden, 1996; La Sorte & Horton, 2021; Lao et al., 2020; Van Doren et al., 2021).

Overall, our analysis reveals that the vast majority of urban bird-building collisions occur on fair weather days, consistent with migration traffic as a major driver. In autumn, collisions also peak on days with high visibility, pointing to the importance of both migration traffic and the appearance of reflections. These immediate predictors of daily collision risk – benign weather, favorable winds, and high visibility at peak migration times – are valuable for predicting days of greatest risk in the future. It is important to note that our study here is limited to urban landscapes. A key question for future work is whether the same patterns occur in residential and suburban sites, which are responsible for a large burden collision mortality across North America (Loss et al., 2014; Machtans et al., 2013). Increased understanding of the conditions and mechanisms that drive bird-building collisions is vital in working towards prevention. As pointed out by Gelb and Delacretaz (2009), there are high-collision sites that are responsible for most of the mortality. New platforms that model and predict migration traffic, such as BirdCast (https://birdcast.info/) and BirdFlow (Fuentes, Van Doren, Fink, & Sheldon, 2022), could be used for focused action (such as turning off lights) on these high-risk sites during the days with highest migration traffic as well as high visibility, yielding the greatest improvements in mitigation measures (Van Doren et al., 2021).

## DATA ACCESSIBILITY

All data and R scripts are available at: https://figshare.com/s/5c455f2a5c763682a737 The repository will be made public when the final version of the study is published.

## AUTHOR CONTRIBUTIONS

RD and KMS designed the study; RD, KMS, PP and AA collected the data; RD and KMS analyzed the data and wrote the manuscript.

## ACKNOWLEDGEMENTS

We are grateful to all of the people who have contributed to the long-term collision monitoring programs at FLAP Canada in Toronto (www.FLAP.org) and Chicago Bird Collision Monitors (www.birdmonitors.net).

