## supplement for "What causes bird-building collision risk? Seasonal dynamics and weather drivers"

**SUPPLEMENTARY MATERIAL**

**Table S1. Models of SPRING collision rates.** Dawn weather was the best predictor of spring collisions (Akaike weight = 0.57). The dawn conditions associated with the greatest collision risk include warm temperatures with a favourable (south) wind direction, low relative humidity, and a lack of precipitation. Predictor variables were centered and standardized to have mean = 0 and SD = 1, except for wind direction which was transformed to range from 0 to 1 and fit with a second-degree orthogonal polynomial. Predictors are indicated in bold where the effect is statistically significant. The sample size is n = 1,465 city-dates. The null model (which includes phenology, but no weather effects) had an Akaike weight of 0.000 (AIC = 3351.1).

|  | MORNING | | |  | **DAWN** | | |
| --- | --- | --- | --- | --- | --- | --- | --- |
|  | AIC = 3221.8 | | |  | **AIC = 3219.9** | | |
|  | Akaike weight = 0.22 | | |  | **Akaike weight = 0.57** | | |
|  | R^2^_GLMM_ = 0.48 | | |  | **R^2^_GLMM_ = 0.48** | | |
| Predictor | Estimate | 95% CI | p | Predictor | Estimate | 95% CI | p |
| **Temperature** | **0.13** | **[0.09, 0.17]** | **< 0.0001** | **Temperature** | **0.14** | **[0.10, 0.18]** | **< 0.0001** |
| Wind speed | –0.01 | [–0.05, 0.04] | 0.80 | Wind speed | –0.02 | [–0.07, 0.02] | 0.27 |
| Poly(Wind direction)-1 | –0.06 | ­[–1.57, 1.45] | 0.93 | Poly(Wind direction)-1 | –0.56 | [–2.13, –1.01] | 0.48 |
| **Poly(Wind direction)-2** | **–4.88** | **[­6.39, –3.37]** | **< 0.0001** | **Poly(Wind direction)-2** | **–4.90** | **[­–6.40, –3.40]** | **< 0.0001** |
| **Precipitation** | **–0.05** | **[–0.14, –0.03]** | **0.02** | **Precipitation** | **–0.05** | **[–0.09, –0.01]** | **0.02** |
| **Relative humidity** | **–0.09** | **[–0.14, –0.03]** | **0.001** | **Relative humidity** | **–0.06** | **[–0.11, –0.02]** | **0.002** |
| Atmos. Pressure | –0.02 | [–0.07, 0.02] | 0.38 | Atmos. Pressure | –0.02 | [–0.06, 0.03] | 0.48 |
| **Cloud cover** | **0.08** | **[0.04, ­0.13]** | **0.0003** | Cloud cover | 0.04 | [0.00, 0.09] | 0.052 |
| Visibility | –0.07 | [–0.12, –0.01] | 0.06 | Visibility | –0.03 | [–0.07, 0.01] | 0.11 |
| Lunar illum. | –0.01 | [–0.05, 0.02] | 0.47 | Lunar illum. | –0.01 | [–0.05, 0.03] | 0.62 |
| **Phenology** | **0.62** | **[0.58, 0.66]** | **< 0.0001** | **Phenology** | **0.62** | **[0.58, 0.66]** | **< 0.0001** |
|  | OVERNIGHT | | |  | PREV. EVENING | | |
|  | AIC = 3232.5 | | |  | AIC = 3221.8 | | |
|  | Akaike weight = 0.003 | | |  | Akaike weight = 0.22 | | |
|  | R^2^_GLMM_ = 0.48 | | |  | R^2^_GLMM_ = 0.48 | | |
|  | Estimate | 95% CI | p |  | Estimate | 95% CI | p |
| **Temperature** | **0.16** | **[0.12, 0.20]** | **< 0.0001** | **Temperature** | **0.14** | **[0.09, 0.18]** | **< 0.0001** |
| Wind speed | –0.02 | [–0.06, 0.02] | 0.29 | Wind speed | ­–0.01 | [–0.05, 0.03] | 0.60 |
| Poly(Wind direction)-1 | 0.12 | [–1.46, ­1.70] | 0.88 | Poly(Wind direction)-1 | –0.77 | [–2.31, 0.77] | 0.33 |
| **Poly(Wind direction)-2** | **–3.42** | **[–4.93, 1.90]** | **< 0.0001** | **Poly(Wind direction)-2** | **–4.47** | **[–6.00, –2.93]** | **< 0.0001** |
| **Precipitation** | **–0.04** | **[–0.08, –0.004]** | **0.03** | Precipitation | 0.00 | [–0.04, 0.04] | 0.96 |
| **Relative humidity** | **–0.07** | **[–0.11, –0.03]** | **0.002** | **Relative humidity** | **–0.10** | **[–0.14, –0.05]** | **< 0.0001** |
| Atmos. Pressure | –0.01 | [–0.05, 0.04] | 0.77 | Atmos. Pressure | ­–0.01 | [–0.06, 0.04] | 0.71 |
| Cloud cover | 0.04 | [–0.01, 0.08] | 0.12 | Cloud cover | 0.02 | [–0.02, 0.07] | 0.32 |
| Visibility | –0.03 | [–0.07, 0.02] | 0.25 | Visibility | –0.03 | [–0.07, 0.02] | 0.22 |
| Lunar illum. | –0.01 | [–0.05, 0.03] | 0.57 | Lunar illum. | –0.02 | [–0.05, 0.02] | 0.36 |
| **Phenology** | **0.62** | **[0.58, 0.66]** | **< 0.0001** | **Phenology** | **0.62** | **[0.59, 0.66]** | **< 0.0001** |

**Table S2. Models of AUTUMN collision rates.** Dawn weather was the best predictor of autumn collisions (Akaike weight > 0.99). The dawn conditions associated with the greatest collision risk include cold temperatures with a favourable (north) wind direction, a lack of precipitation, high visibility, and high atmospheric pressure. Predictor variables were centered and standardized to have mean = 0 and SD = 1, except for wind direction, which was transformed to range from 0 to 1 and fit with a second-degree orthogonal polynomial. Predictors are indicated in bold where the effect was statistically significant. The sample size is n = 1,855 city-dates. The null model (which includes phenology, but no weather effects) had an Akaike weight of 0.000 (AIC = 3670.0).

|  | MORNING | | |  | DAWN | | |
| --- | --- | --- | --- | --- | --- | --- | --- |
|  | AIC = 3393.0 | | |  | AIC = 3332.6 | | |
|  | Akaike weight = 0.00 | | |  | Akaike weight > 0.99 | | |
|  | R^2^_GLMM_ = 0.62 | | |  | R^2^_GLMM_ = 0.61 | | |
| Predictor | Estimate | 95% CI | p | Predictor | Estimate | 95% CI | p |
| **Temperature** | **–0.04** | **[–0.07, 0.00]** | **0.02** | **Temperature** | **–0.05** | **[­–0.08, ­–0.02]** | **0.0006** |
| **Wind speed** | **–0.04** | **[–0.07, –0.01]** | **0.02** | Wind speed | –0.03 | [–0.06, 0.00] | 0.06 |
| **Poly(Wind direction)-1** | **1.70** | **­[0.44, 2.96]** | **0.008** | **Poly(Wind direction)-1** | **3.42** | **[2.17, 4.69]** | **< 0.0001** |
| **Poly(Wind direction)-2** | **4.01** | **[­2.78, 5.24]** | **< 0.0001** | **Poly(Wind direction)-2** | **5.29** | **[­4.06, 6.51]** | **< 0.0001** |
| Precipitation | –0.02 | [–0.05, 0.01] | 0.24 | **Precipitation** | **–0.08** | **[–0.10, –0.05]** | **< 0.0001** |
| Relative humidity | 0.03 | [–0.01, 0.036] | 0.14 | **Relative humidity** | **0.05** | **[0.01, 0.09]** | **0.006** |
| **Atmos. Pressure** | **0.13** | **[0.09, 0.16]** | **< 0.0001** | **Atmos. Pressure** | **0.10** | **[0.06, 0.13]** | **< 0.0001** |
| **Cloud cover** | **–0.05** | **[–0.08, –­0.01]** | **0.004** | Cloud cover | –0.02 | [–0.05, 0.01] | 0.16 |
| **Visibility** | **0.14** | **[0.10, 0.19]** | **< 0.0001** | **Visibility** | **0.14** | **[0.09, 0.18]** | **< 0.0001** |
| **Lunar illum.** | **–0.04** | **[–0.06, –0.01]** | **0.01** | **Lunar illum.** | **–0.03** | **[–0.06, ­–0.01]** | **0.01** |
| **Phenology** | **0.78** | **[0.74, 0.80]** | **< 0.0001** | **Phenology** | **0.78** | **[0.75, 0.81]** | **< 0.0001** |
|  | OVERNIGHT | | |  | PREV. EVENING | | |
|  | AIC = 3400.2 | | |  | AIC = 3353.8 | | |
|  | Akaike weight = 0.00 | | |  | Akaike weight = 0.00 | | |
|  | R^2^_GLMM_ = 0.61 | | |  | R^2^_GLMM_ = 0.62 | | |
|  | Estimate | 95% CI | p |  | Estimate | 95% CI | p |
| **Temperature** | **–0.06** | **[­–0.09, ­–0.02]** | **0.0004** | **Temperature** | **–0.03** | **[–0.06, 0.00]** | **0.03** |
| Wind speed | –0.01 | [–0.05, 0.02] | 0.36 | **Wind speed** | **–0.05** | **[–0.08, –0.02]** | **0.0004** |
| **Poly(Wind direction)-1** | **4.04** | **[2.74, 5.35]** | **< 0.0001** | **Poly(Wind direction)-1** | **5.58** | **­[4.30, 6.86]** | **< 0.0001** |
| **Poly(Wind direction)-2** | **4.71** | **[­3.47, 5.94]** | **< 0.0001** | **Poly(Wind direction)-2** | **7.10** | **[­5.89, 8.32]** | **< 0.0001** |
| **Precipitation** | **–0.04** | **[–0.07, –0.01]** | **0.004** | Precipitation | –0.03 | [–0.05, 0.00] | 0.08 |
| Relative humidity | 0.04 | [0.00, 0.08] | 0.06 | Relative humidity | 0.04 | [0.00, 0.08] | 0.06 |
| **Atmos. Pressure** | **0.08** | **[0.05, 0.12]** | **< 0.0001** | **Atmos. Pressure** | **0.06** | **[0.03, 0.09]** | **0.0003** |
| **Cloud cover** | **–0.06** | **[–0.09, –0.03]** | **0.0003** | **Cloud cover** | **–0.06** | **[–0.09, –­0.02]** | **0.0007** |
| **Visibility** | **0.09** | **[0.04, 0.14]** | **0.0003** | **Visibility** | **0.07** | **[0.02, 0.12]** | **0.004** |
| **Lunar illum.** | **–0.04** | **[–0.06, ­–0.01]** | **0.01** | **Lunar illum.** | **–0.04** | **[–0.06, –0.01]** | **0.007** |
| **Phenology** | **0.78** | **[0.75, 0.81]** | **< 0.0001** | **Phenology** | **0.77** | **[0.74, 0.80]** | **< 0.0001** |
